# Improving polygenic risk prediction in admixed populations by explicitly modeling ancestral-specific effects via GAUDI

**DOI:** 10.1101/2022.10.06.511219

**Authors:** Quan Sun, Bryce T. Rowland, Jiawen Chen, Anna V. Mikhaylova, Christy Avery, Ulrike Peters, Jessica Lundin, Tara Matise, Steve Buyske, Ran Tao, Rasika A. Mathias, Alexander P. Reiner, Paul L. Auer, Nancy J. Cox, Charles Kooperberg, Timothy A. Thornton, Laura M. Raffield, Yun Li

## Abstract

Polygenic risk scores (PRS) have shown successes in clinics, but most PRS methods have focused only on individuals with one primary continental ancestry, thus poorly accommodating recently-admixed individuals. Here, we develop GAUDI, a novel penalized-regression-based method specifically designed for admixed individuals by explicitly modeling ancestry-specific effects and jointly estimating ancestry-shared effects. We demonstrate marked advantages of GAUDI over other methods through comprehensive simulation and real data analyses.

## Main text

Polygenic risk scores (PRS) have been successfully incorporated into clinical risk models for therapeutic interventions and disease screening ^1–3^. However, PRS in personalized medicine disproportionately benefit European ancestry populations ^4^ due to the severe under-representation of non-European ancestry individuals in genetic studies ^5^. Importantly, although multiple methods have been developed for these under-represented populations ^6–12^, few exist to explicitly model information from recently admixed populations. We here present GAUDI, a novel penalized regression based PRS method developed specifically for admixed individuals. GAUDI can model PRS with high accuracy in the presence of ancestry-differentiated effects by balancing fusion and sparsity penalties in a fused lasso ^13^ framework (**Methods**). The fusion component encourages similar effects across ancestries for the same variant, and the sparsity component encourages a small number of variants with non-zero effect.

We choose to model several subsets of variants rather than all genome-wide variants, by applying various thresholds on p-values from standard GWAS, followed by LD clumping to both remove information redundancy and reduce computational burden (**Methods, Extended Data Fig. 1b**). To improve estimation accuracy, we adopt cross-validations to tune parameters, including both the sparsity and fusion parameters and the optimal variant subset (**Methods, Extended Data Fig. 1c**). Finally, we calculate PRS for each target individual with the estimated effect size parameters.

We evaluated the performance of GAUDI through comprehensive simulations. We first compared GAUDI with the clumping and thresholding method implemented in PRSice ^14^ and the previously proposed partial PRS (pPRS) ^15^ method, under the scenario of no ancestry-specific effects (*p*_shared_ = 1, **Methods**). While 100% of effects being shared across ancestry is an over-simplification, recent work has shown that there is almost always a positive correlation between effect sizes across global populations for most variants associated with complex traits ^16^. We ran PRSice, pPRS, GAUDI with and without LD clumping for comparison (**Methods, Supplementary Notes**). We used COSI ^17^ to simulate 500kb regions for 3,500 AA individuals assuming 80% African (AFR) and 20% European (EUR) admixture, and another independent samples of 2,500 EUR and 2,500 AFR individuals serving as references. We considered three different genetic settings of the causal variants in terms of their minor allele frequency (MAF) across ancestries: variants with EUR-MAF and AFR-MAF both >= 5% (causal variants common in both ancestries), variants with EUR-MAF >= 5% and AFR-MAF < 5% (casual variants common only in EUR), and variants with EUR-MAF < 5% and AFR-MAF >= 5% (causal variants common only in AFR). For each of three MAF settings, we varied the proportion of causal variants to be 1, 0.5, 0.05 to represent different polygenicity situations, and the proportion of variation explained by genetic variations (i.e., heritability) to be 0.2 or 0.6. In addition, we also varied the maximum LD R^2^ among causal variants to be 0.2 or 0.5.

Under all the three cross-ancestry MAF settings, GAUDI outperformed PRSice and pPRS across all simulated traits in the held-out testing data (**Fig. 1 a-c, Extended Data Fig. 2 a-c, Supplementary Table 1, Supplementary Notes**). Comparing across different polygenicity and heritability scenarios, GAUDI achieved best performance across the entire spectrum assessed, demonstrating most pronounced performance gains in settings with higher heritability and denser genetic architecture. In addition, the R^2^ attained by GAUDI in the testing dataset is nearly equal to heritability in almost all simulated phenotypes, demonstrating the power of GAUDI by borrowing information from haplotype segments in one ancestry to better estimate the effects in another ancestry.

**Fig 1.**
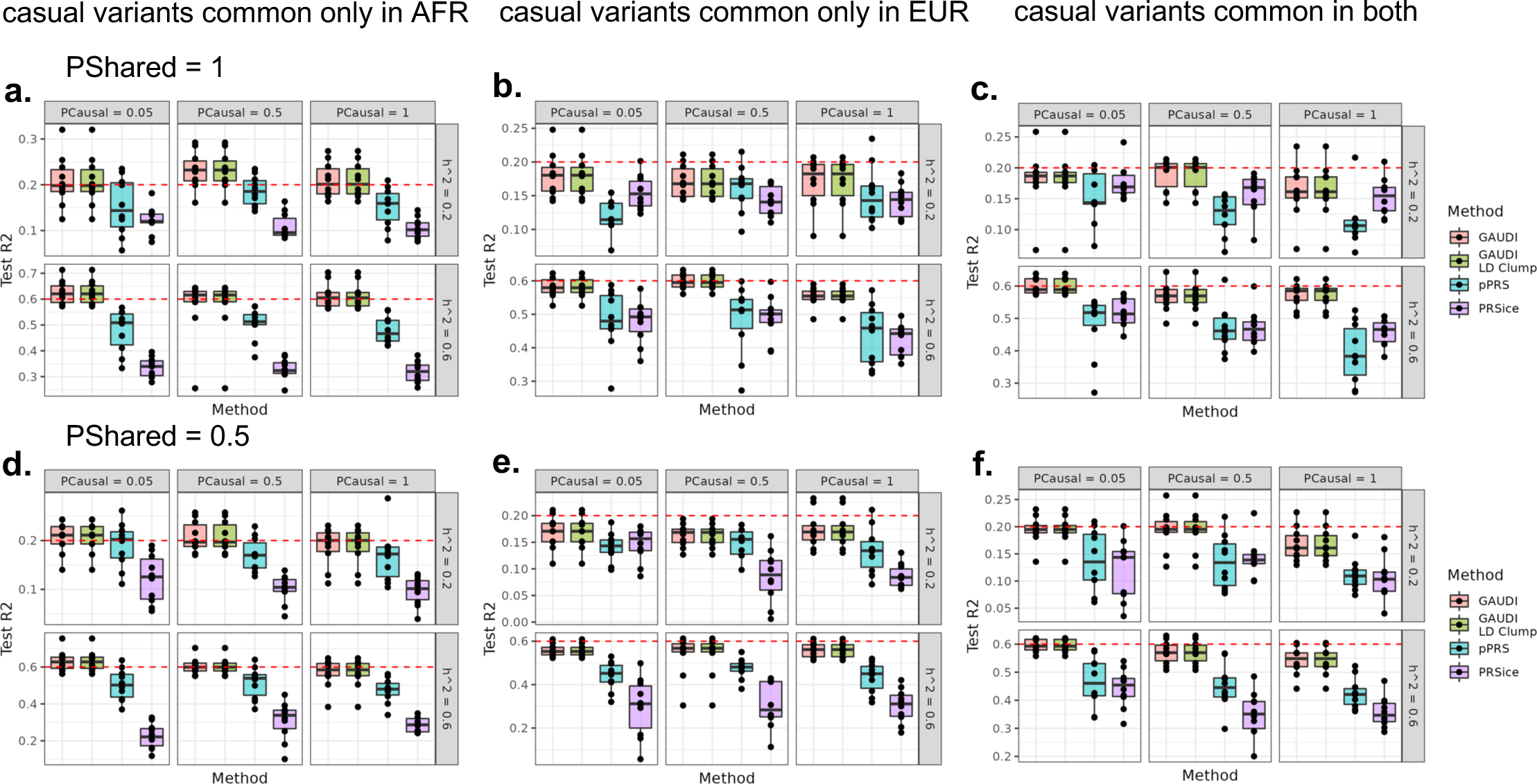
GAUDI performance compared to PRSice and pPRS in simulation studies under different settings. (a)-(c). PShare (proportion of variants with shared effects across ancestry groups) = 1: no ancestry-specific effects for all causal variants. **(d)-(e)**. PShare = 0.5: half of the causal variants have ancestry-specific effects. **(a)(d)**. Causal variants are common only in AFR ancestry, specifically EUR-MAF < 5% and AFR-MAF >= 5%. **(b)(e)**. Causal variants are common only in EUR ancestry, i.e., EUR-MAF >= 5% and AFR-MAF < 5%. **(c)(f)**. Causal variants are common in both ancestries, i.e., EUR-MAF and AFR-MAF both >= 5%. Each experiment was repeated 10 times. The maximum LD R2 between causal variants were set to be 0.2 for all settings. The dashed red line denotes heritability. PCausal: proportion of causal variants out of all variants.

We then simulated phenotypes where 50% of causal variants have ancestry-specific effects and the remaining 50% have effect sizes shared across the two ancestral populations (**Methods**). Similarly, we considered multiple genetic architectures by varying causal variants’ cross-ancestry MAFs, heritability, polygenicity and maximum LD R^2^. Our results were largely consistent with those from the above simulations (**Fig. 1 d-f, Extended Data Fig. 2 d-f, Supplementary Table 2**). The improvement of GAUDI over competing methods is even more pronounced in some scenarios with the introduction of ancestry-specific effects (**Fig. 1**). We also note the variability of GAUDI is slightly reduced compared to the previous setting where no ancestry-specific effects were allowed. These results further underscore the advantage of GAUDI by allowing and jointly modeling ancestry-specific effects. We also simulated phenotypes where the proportion of ancestry-specific causal variants is 20%, and the results are highly consistent (**Extended Data Fig. 3**), suggesting GAUDI is robust to a variety of genetic architectures. Furthermore, GAUDI with LD clumping performs almost identically well as GAUDI without LD clumping, indicating GAUDI is also robust to the inclusion of correlated variants in the PRS construction process.

We then performed real data analysis for African American (AA) from the Women’s Health Initiative (WHI) study (**Methods, Supplementary Notes**). We added PRS-CSx in our method comparison given its popularity in recent PRS literature ^6–12^, along with PRSice and pPRS. We considered five phenotypes including white blood cell count (WBC), platelet count (PLT), hematocrit (HCT), hemoglobin (HGB), and C-reactive protein (CRP). Across the five phenotypes, only CRP and WBC showed significant non-zero mean R^2^ (**Fig. 2, Supplementary Table 3**). The relative order and magnitude of prediction accuracy we observed in the WHI AA samples are expected for HCT, HGB and PLT, consistent with recent applications of PRS to blood cell traits in AA samples ^18^. For CRP and WBC, GAUDI substantially improves prediction accuracy compared to alternative methods. For example, GAUDI could achieve testing R^2^ of 1% - 3%, while the other three methods result in almost negligible R^2^ (<1%). The relative improvement in the mean R^2^ by GAUDI is 63.8% and 406.4% compared to PRS-CSx, 93.4% and 567.6% compared to PRSice, and 169.7% and 758.7% compared to pPRS, for WBC and CRP respectively. Such improvements are striking especially given the fact that GAUDI only utilizes variants with AFR GWAS p-value < 5e-5, while PRS-CSx and PRSice considered all variants evaluated in GWAS. For example, GAUDI modeled an average of only 65 variants across the five folds to construct WBC PRS, while PRS-CSx and PRSice used >500,000 variants. These results demonstrate the advantage of GAUDI by allowing differential effects across ancestry, suggesting that explicit modeling of the genetic mosaicism in recently admixed populations can be much more rewarding and influential than simply including more variants in PRS construction.

**Fig 2.**
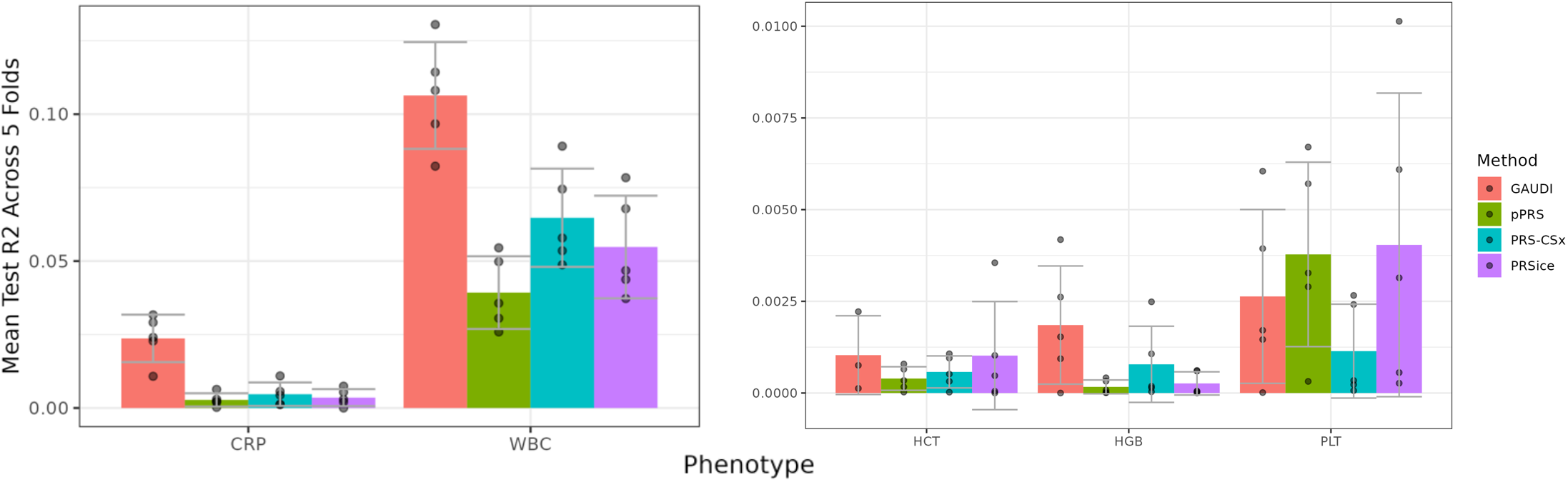
GAUDI performance compared to pPRS, PRS-CSx and PRSice in WHI AA samples. Each analysis was repeated five times, with different training and testing samples. The error bar denotes the standard error across replicates. CRP: C-reactive protein, WBC: white blood cell count, HCT: hematocrit, HGB: hemoglobin, PLT: platelet count.

While CRP and WBC are extreme examples where large ancestry-specific effects exist, the ability of GAUDI to capture such extremes is clinically meaningful. A recently publication shows that the Duffy null variant (rs2814778) should be accounted for in clinical decision-making to avoid unnecessary bone marrow biopsy procedures ^19^. Furthermore, our simulation results demonstrate that GAUDI still performs better than other methods even if there are no extreme large ancestry-specific effects. In addition, the real data results for HCT, HGB and PLT show that GAUDI achieves similar or better performance over other methods for traits without known large ancestry-specific effects.

In summary, both comprehensive simulations and real data analysis demonstrate the superiority of GAUDI over alternative methods by allowing ancestry-specific effects, which we anticipate will be increasingly observed with larger numbers of non-European ancestry individuals evaluated in genetic association studies. We note that GAUDI utilizes individual level data, which is more demanding computationally and data-wise than methods based on summary statistics, which could be addressed in future research. We believe with more admixed individuals enrolled in more studies in the coming years, the community will benefit even more from GAUDI, particularly to avoid further exacerbating health disparity for admixed individuals.

## Methods

### Model setup

Consider the problem of constructing PRS for a sample of *i =* 1, …, *n* individuals recently admixed from two ancestral populations, *A* and *B*. This model can be extended to an arbitrary number of ancestral populations, but for simplicity here we consider only two ancestral populations. Let *x*_*i j* 1_, *x*_*i j* 2_ denote the allelic value of individual *i* for variant *j* on haplotype 1 and 2, respectively (**Extended Data Fig. 1a**), taking values 0 or 1 for genotype data, or ranging continuously from 0 to 1 for imputed dosages. Similarly, let *l*_*i j* 1_, *l*_*i j* 2_ denote the local ancestry for individual *i* for variant *j* on haplotype 1 and 2, respectively, taking values A or B for the corresponding ancestral population. Let *Y* = (*y*_1_, … *y*_*n*_)′ be an n × 1phenotype vector, and we assume

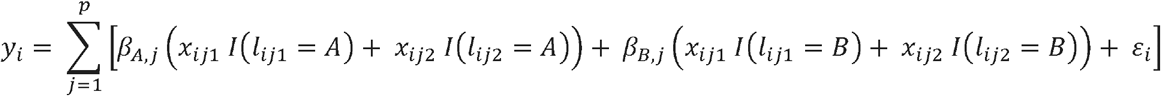

where *p* is the total number of variants, and *I* (·) is the indicator function. A subset of the variants, *p*\*, are causal, meaning that the effect of the variants on the phenotype is non-zero. Under this model, *β*_*A, j*_, *β*_*B, j*_ are the population *A, B* specific effect of variant *j* on the phenotype. With no local ancestry information, nor regards to haplotype information, this collapses to the usual genetic association model

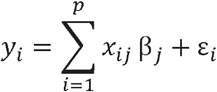

Where *x*_*ij*_ is the allelic values of individual *i* for variant *j*, and *β* _*j*_ is the effect size of variant *j*.

We further write the above model using matrix notation. Let *x*_*i j P*_ = *x*_*i j* 1_ *I (l*_*i j* 1_ = *P)* + *x*_*i j* 2_ (*l*_*i j* 2_ = *P*) where *P* denotes population ancestry, taking values *A* or *B*. Then, the design matrix is given by

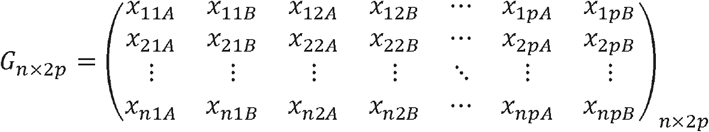

We then define *β*_2*p* × 1_ = (*β*_*A*,1_, *β*_*B*,1_, … *β*_*A, p*_, *β*_*B,p*_), thus the above phenotype model could be represented as, *Y* = *G* β + ε, where *ε* = (*ε*_1_, …, *ε*_*n*_) is the error vector.

The problem of PRS construction under this model is equivalent to the problem of accurate estimation of the population specific effects for ancestral populations *A* and *B* with the design matrix specified, and given that the predictors (variants) are already selected.

### GAUDI framework

Our GAUDI method for PRS construction for admixed individuals is a modified fused lasso approach. Specifically, given genotype information for *n* admixed individuals at *p* variants, some subset of which (denoted by *p*\*) are causal variants. We assume that for each individual we have also obtained haplotype-resolved local ancestry inference estimates via RFMix.

The variant selection workflow is shown in **Extended Data Fig. 1b**. Using the training sample, we perform GWAS and select variants based on GWAS p-values. Note that it is also acceptable to use external GWAS results to select variants, which can be preferred for at least two reasons. First, we can leverage information from larger sample size and thus more powerful GWAS already carried out. Second, using external GWAS results will save computation costs for running GWAS in the training sample. With GWAS p-values, we adopt a grid search strategy to select variants. For *k* pre-specified *p*-value thresholds, (*t*_1_, …, *t*_*k*_), we can identify *k* sets of variants passing each of the thresholds. Then we perform LD clumping on each of the *k* selected variant sets to both reduce dimension and remove variants in high LD for more stable inference. Let *p*_*t*_ denote the total number of variants for the set of variants selected with *p*-value threshold *t*. We then adopt a grid search strategy via five-fold cross validation to estimate the best tuning parameters using the following fused lasso objective function :

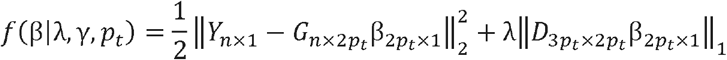

where the penalty matrix *D* is given by

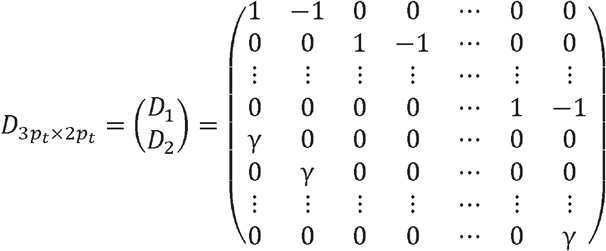

Then we compare the optimized performance for *k* variant sets with different p-value thresholds, and report the best one as the final constructed PRS model (**Extended Data Figure 1c**).

One notable difference between GAUDI and traditional fused lasso is that only ancestry-specific effects for a given variant are penalized with fusion, rather than all adjacent parameters. We finally calculate the PRS for a target sample using

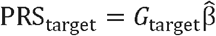

Cross-validated model performance for tuning parameters (λ, γ and the *p*-value threshold *t*_*i*_) is optimized based on the squared Pearson correlation between the observed phenotype and the PRS calculated above.

### COSI genotype simulations

In order to simulate haplotypes of recent admixture, we used COSI ^17^ to generate 500kb regions for 3,500 AA individuals. We made two primary assumptions in generating our simulated haplotypes. First, we assumed that the global ancestry proportions of our AA samples were 80% African and 20% European. Second, using empirical estimates of ancestral switch-points based on an analysis of TOPMed individuals ^20^, we assumed 4% of 500Kb regions would contain ancestry switch-point events. Thus, for 3,500 diploid individuals, 280 chromosomes contained switch points (7,000*0.04 = 280). For each ancestry switch point chromosome, we generated one EUR and one AFR chromosome to simulate the admixture event at a random base-pair in the region. For the remaining 6,720 chromosomes with no admixture events, we generated 80% AFR chromosomes (n = 5,376) and 20% European chromosomes (n = 1,344). Additionally, we simulated 5,000 EUR chromosomes and 5,000 AFR chromosomes to be used as reference for relevant methods.

### Phenotype simulations

We simulated phenotypes using 500kb regions generated from COSI simulated genotypes for the 3,500 admixed individuals and the 2,500 reference AFR and EUR individuals. We considered three distinct sets of causal variants to mimic different genetic architectures.

First, we created the “causal variants common in both ancestries” scenario. At a locus, we considered variants that had both AFR MAF and EUR MAF >= 0.05 as candidate causal variants. Second, we created the “causal variants common only in EUR” scenario, where variants that had AFR MAF < 0.05 and EUR MAF >= 0.05 were considered as candidate causal variants. Third, we similarly created the “causal variants common only in AFR” scenario, where variants that had AFR MAF >= 0.05 and EUR MAF < 0.05 were considered as candidate causal variants.

For a variant *j*, we simulated its effect the sizes from the following distribution

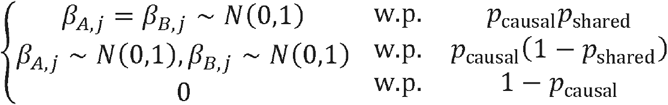

We changed the values of four different parameters to evaluate a wide spectrum of genetic architectures. First, we varied the proportion of causal variants (*p*_casual_), taking three possible values 0.05, 0.5 and 1, to represent different levels of polygenicity. Second, we varied the proportion of variants that have the same effect size across ancestry groups (*p*_shared_), taking three possible values 1, 0.8, 0.5, to represent varying extents of genetic heterogeneity across ancestries. Third, we varied heritability (*h*^2^), or the proportion of variation explained by genetic effects, taking possible values 0.2 or 0.6. Finally, we allowed different levels of maximum correlation between causal variants (*r*^2^), up to 0.2 and 0.5, to test GAUDI model stability in the presence of correlated causal variants.

For varying the LD between causal variants in the phenotype, we performed LD pruning on the set of candidate causal variants using PLINK (–indep-pairwise 500 5 *r*^2^)^21^. We repeated each combination of the above parameters 10 times for each of the three causal variant scenarios. We controlled the total by estimating the variance explained by the causal variants, and then heritability at a desired *h*^2^ simulating error terms from a normal distribution with a particular standard deviation.

### The WHI cohort

The Women’s Health Initiative (WHI) is one of the largest (n=161,808) studies of women’s health ever undertaken in the U.S. There are two major components of WHI: (1) a clinical trial (CT) that enrolled and randomized 68,132 women ages 50-79 into at least one of three placebo control clinical trials (hormone therapy, dietary modification, and supplementation with calcium and vitamin D); and (2) an observational study (OS) that enrolled 93,676 women of the same age range into a parallel prospective cohort study ^22^. A diverse population including 26,045 (17%) women from minority groups was recruited from 1993-1998 at 40 clinical centers across the U.S. Details on the study design, eligibility, recruitment, and the reliability of the baseline measures of demographic and health characteristics have been published elsewhere ^22,23^ Fasting blood samples were obtained from all participants at baseline and were analyzed for white blood cell count and platelet count by certified laboratories at each of the 40 clinical centers as part of a complete blood count (CBC) ^23^. Results were entered into the WHI database at each clinical center and were reviewed by clinical center staff ^24^. These assays were performed in a single laboratory using the same methods. CBCs were measured within 30 hours of blood draw.

The **WHI PAGE GWAS** project performed genotyping among self-identified non-Hispanic Black or African American (n=6897) and Hispanic/Latino (n=4754) women from WHI who consented to genetic research. These participants were genotyped by the Population Architecture using Genomics and Epidemiology (PAGE) study, along with participants of non-European ancestry from the Hispanic Community Health Study/Study of Latinos (HCHS/SOL), the Multiethnic Cohort (MEC), and the Icahn School of Medicine at Mount Sinai BioMe biobank (BioMe) (total MEGA sample size n=49,839). Genotyping was performed using the Multi-Ethnic Genotyping Array (MEGA); quality control has been described previously ^25^ and included exclusion of variants based on high missingness, Mendelian error rates, discordant calls among study duplicate samples, and other filters. This array was designed to provide improved multi-ethnic coverage of common and low frequency variants, and also included custom content for fine-mapping GWAS loci and genotyping clinically reported and exonic variants ^26^.

The **WHI WHIMS+ GWAS** project performed genotyping among women of European descent with appropriate consent for genetic data sharing on dbGaP using the Illumina Omni Express platform. When these participants are combined with the GARNET (Genomics and Randomized Trials Network from NHGRI) participants (who were genotyped on the Illumina Omni-Quad chip), they form a population that is representative of the entire European American hormone trial population from WHI.

### Real data analysis on WHI samples

In this study, we included 6,734 AA individuals from the WHI PAGE GWAS study and 5,681 EUR individuals from WHI WHIMS+ GWAS study, to compare the performance of GAUDI with PRSice, pPRS, and PRS-CSx. The EUR individuals were included as ancillary samples in order to apply pPRS and PRS-CSx, both of which require EUR GWAS estimates as input. We used 5-fold cross validation to assess performance of different methods.

#### Genotype imputation

Genotype imputation was performed with TOPMed freeze 8 reference panel ^27^ following the procedure of our previous work ^28–31^, using Eagle v2.4 ^32^ for phasing and minimac4 ^33^ for imputation. We performed imputation separately for AA samples or EUR samples. Starting from the genotype array data, we removed samples and variants with missingness > 10%, and then uploaded the data to TOPMed imputation server to perform imputation. After imputation, we re-calculated the estimated imputation quality (Rsq) to account for sample overlap with the reference panel, and performed post-imputation QC by including well-imputed variants with imputation Rsq > 0.3 for common variants (MAF > 1%) and imputation Rsq > 0.6 for low frequency variants (MAF in [0.1%, 1%]).

#### Phenotype QC

We considered four blood cell phenotypes (WBC, PLT HCT, HGB) and CRP, all five traits with low levels of missing data. All phenotypes were adjusted by cohort for age, squared age, top 10 genotype PCs, recruitment center and genotyping array using linear regression models. WBC values were log10(x + 1) transformed before regression. Residuals from the regression models were inverse normal transformed and served as the phenotypes for GWAS analysis.

#### GWAS

For the GWAS association tests, we considered common variants (MAF > 0.01) with Rsq > 0.3, and low frequency variants (MAF in (0.001, 0.01)) with Rsq > 0.6. Note that for our training samples, MAF = 0.001 corresponds to a minor allele count (MAC) of approximately 10. We performed GWAS using REGENIE ^34^ separately for each of the five training sets of AA individuals (i.e., for 5-fold CV), and for all the EUR individuals, on the residuals of each phenotype obtained in linear regression models adjusting for age, age squared, top 10 genotype PCs, recruitment center and genotyping array. To fit the REGENIE null model accounting for cryptic relatedness, we used extremely-well imputed common variants (MAF > 0.2, Rsq > 0.9999). We fit the five phenotypes simultaneously using the grouping options available in REGENIE with default parameters.

### Local ancestry inference

For the AA samples in both simulations and real data analysis for the WHI participants, we inferred local ancestry using RFMix ^35^ with data from the 1000 Genomes Project (1000G) ^36^ as the reference panel. We considered only EUR and AFR ancestry since our analyses focused on AAs. Specifically, our 1000G reference panel included 92 EUR samples and 92 and AFR samples. For local ancestry inference, we kept only common variants with MAF > 0.05.

### PRS method application

#### GAUDI

When applying GAUDI, we included variants that had a MAC > 10 on at least one ancestral haplotype. If the variant was polymorphic in only one ancestral population, we included only one ancestry-specific effect in the model. If the variant was polymorphic in both populations, we included both ancestry-specific effects in the model. Other details of GAUDI were described previously in the **GAUDI framework** section.

#### PRSice

PRSice is a popular software implementation of the P+T or C+T method ^14^. We applied PRSice to the REGENIE summary statistics in both AA individuals and EUR individuals without using local ancestry information. We ran PRSice on training samples with default parameters, and we then applied the formula (PRSice selected variants and their weights estimated from training samples) on testing samples to obtain the weighted sum, which was the PRS for testing samples.

#### Partial PRS (pPRS)

pPRS is a method to incorporate local ancestry information in PRS estimation in admixed individuals using only summary statistics from the ancestral populations ^15^. We applied pPRS with default parameters using GWAS results from EUR and AFR reference samples, and using RFMix estimated local ancestry for each target sample.

#### PRS-CSx

PRS-CSx is a recently developed method that integrates GWAS summary statistics from multiple populations while accounting for LD from external reference panels to improve cross-population PRS prediction ^12^. We applied PRS-CSx with both AA and EUR GWAS summary statistics without using local ancestry information.

For all the methods, PRS performance was assessed by mean testing R2 between PRS and adjusted phenotypes across multiple repeats.

## Acknowledgement

This study was supported by NIH grant U01HG011720 and R01HL151152. In addition, LMR was also supported by R01HG010297 and by the National Center for Advancing Translational Sciences, National Institutes of Health, through Grant KL2TR002490. QS was also supported by U24AR076730. The WHI program is funded by the National Heart, Lung, and Blood Institute, National Institutes of Health, U.S. Department of Health and Human Services through contracts 75N92021D00001, 75N92021D00002, 75N92021D00003, 75N92021D00004, 75N92021D00005.

## Data and Codes Availability

WHI data could be accessed through dbGaP at phs000200 or upon application to the WHI Coordinating Center (https://www.whi.org/).

Codes used for analyses are available at https://github.com/brycerowland/GAUDI.

## Author Contributions

Study design and conceptualization: QS, BTR, AVM, RT, RAM, APR, PLA, NJC, TAT, LMR, YL; analysis: QS, BTR, JC; data generating and coordinating: CA, UP, JL, TM, SB, LMR; manuscript writing: QS, BTR; supervision: YL. All authors contributed to manuscript revisions.

