## Extended Data Figures for "Improving polygenic risk prediction in admixed populations by explicitly modeling ancestral-specific effects via GAUDI"

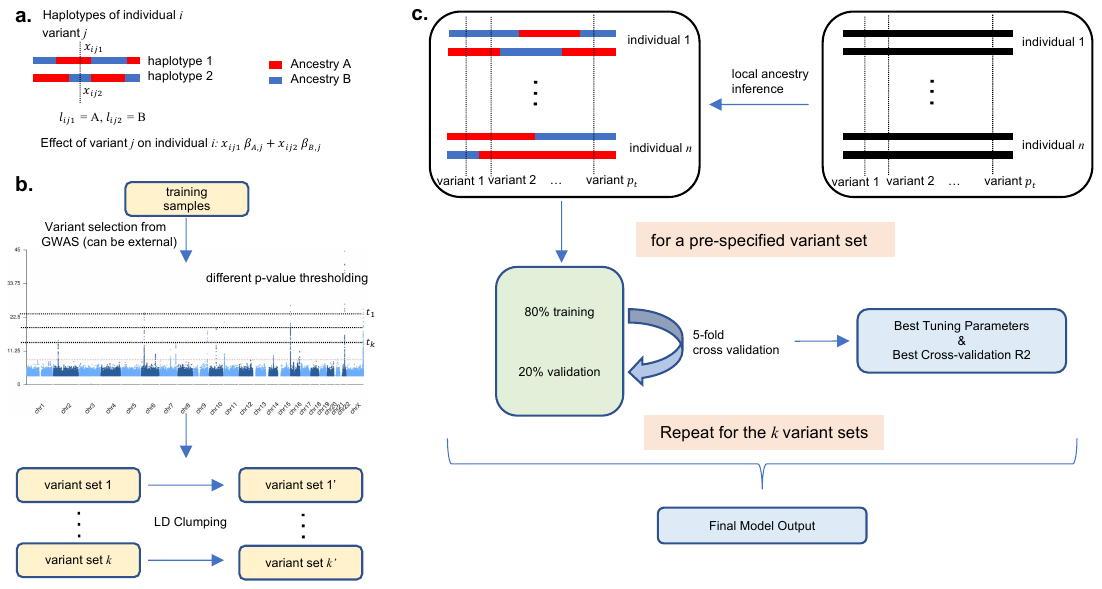

**Extended Data Fig 1. Overview of GAUDI model and framework. (a) Model set-up of GAUDI.** Consider the haplotypes of individual *i* at variant *j* and suppose local ancestry is already inferred. We consider the scenario of only two ancestries, namely A and B. Let $x_{ij1}, x_{ij2}$ denote the haplotype values (taking values 0 or 1 for a directly genotyped variant, and ranging from 0 to 1 for an imputed variant). Let $l_{ij1}, l_{ij2}$ denote the local ancestry; here we have $l_{ij1}=A, l_{ij2}=B$. Let $\beta_{A,j},\beta_{B,j}$ denote the population A, B specific effect of this variant on the phenotype. Thus we have the total effect of variant *j* in individual *i* as $x_{ij1}\beta_{A,j}+x_{ij2}\beta_{B,j}$. **(b) Variant selection framework of GAUDI.** We first perform GWAS or use external GWAS results to obtain p-values, which will be used for variant selection. Specifically, we use the thresholding strategy to identify variants that are marginally associated with the trait of interest at *k* pre-specified *p*-value thresholds, $\left( t_{1},\cdots,t_{k} \right)$. These *k* sets of variants will be generated, and we then perform LD clumping for each of the *k* sets to both reduce dimension and remove variants in high LD. **(c). Final PRS construction of GAUDI.** After inferring the local ancestry for every participant in the training set, for a specific set of $p_{t}$ variants, we split the training individuals into 80% training and 20% testing and perform five-fold cross-validation to select the best tuning parameters, under the penalized regression framework. Repeating the process for the *k* variant sets and comparing the cross-validated R2 will give us the final PRS model.

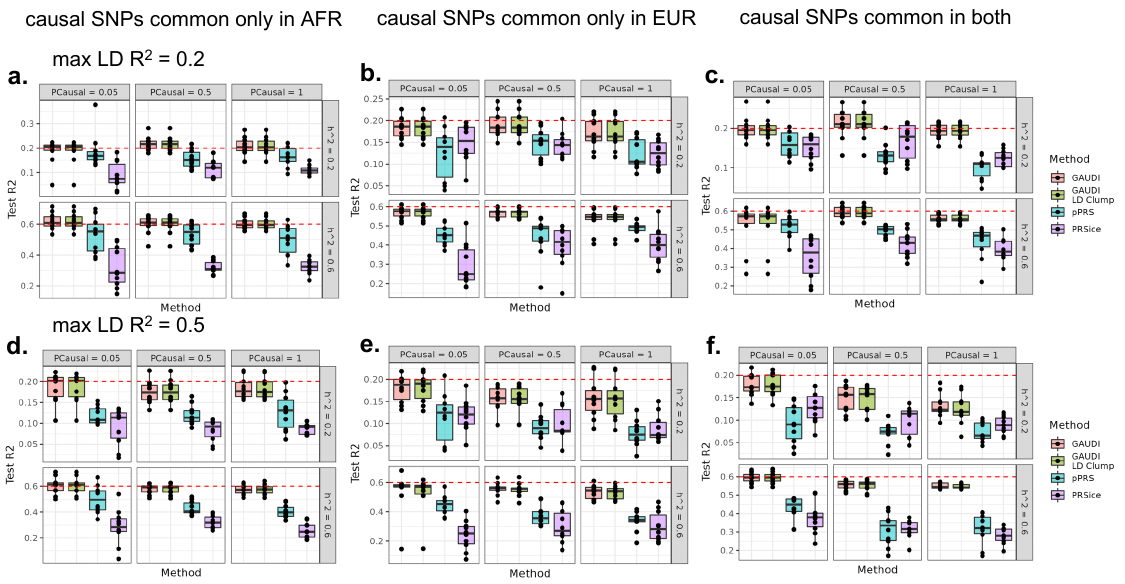

**Extended Data Fig 2. GAUDI performance compared to PRSice and pPRS in simulation studies under different settings, with maximum LD R2 between causal variants to be 0.5. (a)-(c).** PShare (proportion of variants with shared effects across ancestry groups) = 1: no ancestry-specific effects for all causal variants. **(d)-(e).** PShare = 0.5: half of the causal variants have ancestry-specific effects. **(a)(d).** Causal variants are common only in AFR ancestry, specifically EUR-MAF < 5% and AFR-MAF >= 5%. **(b)(e).** Causal variants are common only in EUR ancestry, i.e., EUR-MAF >= 5% and AFR-MAF < 5%. **(c)(f).** Causal variants are common in both ancestries, i.e., EUR-MAF and AFR-MAF both >= 5%. Each experiment was repeated 10 times. The maximum LD R2 between causal variants were set to be 0.5 for all the settings. The dashed red line denotes heritability. PCausal: proportion of causal variants out of all variants.

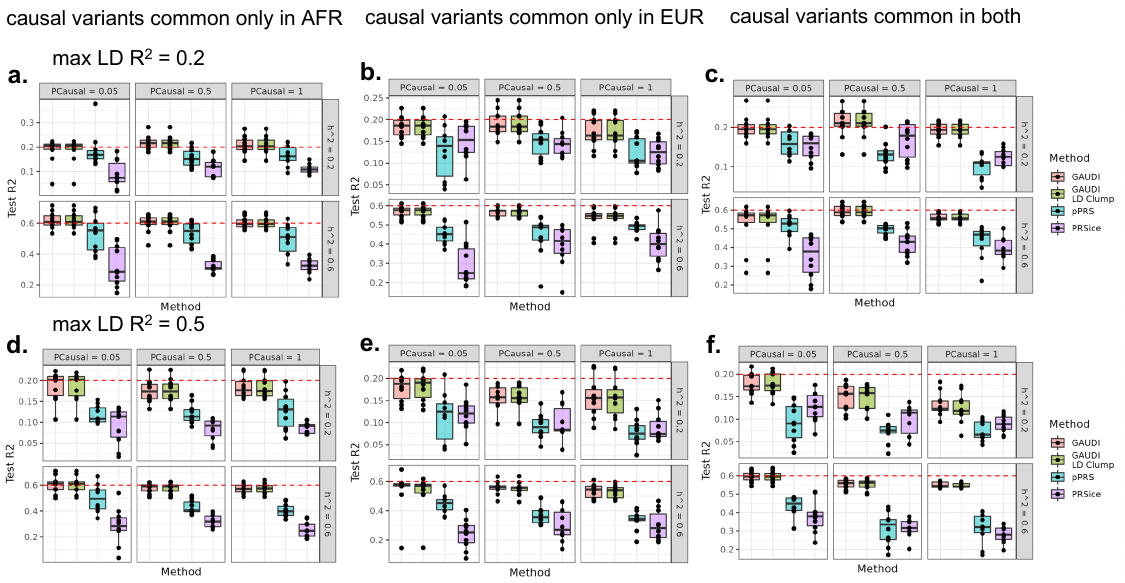

**Extended Data Fig 3.** **GAUDI performance compared to PRSice and pPRS in simulation studies under different settings, with the proportion of ancestry-specific causal variants being 20%.** **(a)-(c).** The maximum LD R^2^ between causal variants were set to be 0.2. **(d)-(e).** The maximum LD R^2^ between causal variants were set to be 0.5. **(a)(d).** Causal variants are common only in AFR ancestry, specifically EUR-MAF < 5% and AFR-MAF >= 5%. **(b)(e).** Causal variants are common only in EUR ancestry, i.e., EUR-MAF >= 5% and AFR-MAF < 5%. **(c)(f).** Causal variants are common in both ancestries, i.e., EUR-MAF and AFR-MAF both >= 5%. Each experiment was repeated 10 times. The proportion of ancestry-specific causal variants was set to be 20% for all the settings. The dashed red line denotes heritability.
