## Supplementary Notes for "Improving polygenic risk prediction in admixed populations by explicitly modeling ancestral-specific effects via GAUDI"

**Simulation results under no ancestry-specific effects**

For the 3,500 AA individuals, we inferred local ancestry with RFMix [^26^](https://sciwheel.com/work/citation?ids=3777659&pre=&suf=&sa=0&dbf=0) using haplotypes from the 1000 Genomes Project [^16^](https://sciwheel.com/work/citation?ids=790619&pre=&suf=&sa=0&dbf=0) as reference, assuming two-way admixture between EUR and AFR. In parallel, we performed GWAS with REGENIE [^27^](https://sciwheel.com/work/citation?ids=11074244&pre=&suf=&sa=0&dbf=0) separately for the 2,500 EUR and AFR reference individuals. When applying GAUDI, we adopted a variant selection strategy to allow for ancestry-specific effects (**Methods, Extended Data Fig. 1b**). For each of the 3,500 target individuals, we constructed PRSice PRS with GWAS results from the 2,500 AFR reference individuals. To obtain pPRS for the target individuals, we applied a piecewise combination of AFR and EUR GWAS weights based on the inferred local ancestry information. We then calculated the squared Pearson correlation (R2) between each of the PRSs (PRSice, pPRS, GAUDI without LD pruning, and GAUDI with LD pruning) and phenotype in 3,500 AA testing samples, which served as the quality metric to compare across methods. We repeated simulations 10 times for each scenario to assess performance variability.

Under the setting where causal SNPs are common only in AFR, we observed that pPRS tends to outperform PRSice, as expected, by accounting for local ancestry. But the improvement is less impressive compared to GAUDI (**Fig. 1a**). For the setting where causal SNPs are common only in EUR, where phenotypes reflect the current bias toward common EUR variants in published GWAS results, PRSice performs better than pPRS in the low heritability scenario, but slightly worse than pPRS in the high heritability scenario (**Fig. 1b**). Despite that EUR segments comprise only 20% of the simulated haplotypes on average, GAUDI demonstrated substantial gains by borrowing information across ancestral populations to construct PRS. The substantial increase in predictive performance of GAUDI over alternative methods suggests an immediate improvement by applying GAUDI to construct PRS in admixed individuals, even when using previously published GWAS variants that are discovered in large European cohorts. Under the setting where causal SNPs are common in both ancestries, pPRS is inferior to PRSice in most cases (**Fig. 1c**). The performance comparison between pPRS and PRSice suggests the limitation of simply combining ancestry-specific PRS when lacking enough AFR common causal variants in input GWAS studies. In contrast, GAUDI still shows substantial improvement. These results demonstrate the utility of GAUDI across a range of genetic architectures in admixed populations.

**Simulation results with ancestry-specific effects**

We observed that GAUDI PRS captured almost all the heritability in testing samples for most phenotypes under the "causal SNPs common only in AFR'' setting (**Fig. 1d**). It also achieved considerable gains relative to alternative methods in the settings of "causal SNPs common only in EUR" (**Fig. 1e**) and "causal SNPs common in both ancestries” (**Fig. 1f**). Under the simulation framework where causal SNPs are common in both ancestries, the improvement of GAUDI over PRSice is much larger when ancestry-specific effects are introduced (**Fig. 1f**) compared to simulated phenotypes without ancestry-specific effects (**Fig. 1c**). These results demonstrate that GAUDI can accurately estimate both ancestry-shared and ancestry-specific effects to construct better PRS in admixed individuals.
